# Development of CpG-adjuvanted stable prefusion SARS-CoV-2 spike antigen as a subunit vaccine against COVID-19

**DOI:** 10.1101/2020.08.11.245704

**Authors:** Tsun-Yung Kuo, Meei-Yun Lin, Robert L Coffman, John D Campbell, Paula Traquina, Yi-Jiun Lin, Luke Tzu-Chi Liu, Jinyi Cheng, Yu-Chi Wu, Chung-Chin Wu, Wei-Hsuan Tang, Chung-Guei Huang, Kuo-Chien Tsao, Shin-Ru Shih, Charles Chen

## Abstract

The COVID-19 pandemic caused by the novel coronavirus SARS-CoV-2 is a worldwide health emergency. The immense damage done to public health and economies has prompted a global race for cures and vaccines. In developing a COVID-19 vaccine, we applied technology previously used for MERS-CoV to produce a prefusion-stabilized SARS-CoV-2 spike protein by adding two proline substitutions at the top of the central helix (S-2P). To enhance immunogenicity and mitigate the potential vaccine-induced immunopathology, CpG 1018, a Th1-biasing synthetic toll-like receptor 9 (TLR9) agonist was selected as an adjuvant candidate. S-2P was combined with various adjuvants, including CpG 1018, and administered to mice to test its effectiveness in eliciting anti-SARS-CoV-2 neutralizing antibodies. S-2P in combination with CpG 1018 and aluminum hydroxide (alum) was found to be the most potent immunogen and induced high titer of spike-specific antibodies in sera of immunized mice. The neutralizing abilities in pseudotyped lentivirus reporter or live wild-type SARS-CoV-2 were measured with reciprocal inhibiting dilution (ID_50_) titers of 5120 and 2560, respectively. In addition, the antibodies elicited were able to cross-neutralize pseudovirus containing the spike protein of the D614G variant, indicating the potential for broad spectrum protection. A marked Th-1 dominant response was noted from cytokines secreted by splenocytes of mice immunized with CpG 1018 and alum. No vaccine-related serious adverse effects were found in the dose-ranging study in rats administered single- or two-dose regimens with up to 50 μg of S-2P combined with CpG 1018 alone or CpG 1018 with alum. These data support continued development of CHO-derived S-2P formulated with CpG 1018/alum as a candidate vaccine to prevent COVID-19 disease.

## Introduction

COVID-19 was first identified as a cause of severe pneumonia cases in December 2019 in association with a seafood market in Wuhan, China [1]. The viral agent was identified as a novel SARS-like coronavirus (SARS-CoV-2) most closely related to bat coronavirus [1]. In the six months since its first appearance, SARS-CoV-2 has become the largest pandemic since the 1918 influenza with nearly 20 million infected and over 700,000 deaths worldwide as of August 2020 [2, 3]. The rapid spread and huge socioeconomic impact of this pandemic require the urgent development of effective countermeasures, including vaccines. In response, pharmaceuticals, academia, and institutions are developing vaccines and drugs at an unprecedented pace with governments and foundations pledging hundreds of millions of dollars for COVID-19 research [4].

In addition to basic public health control measures such as social distancing, contact tracing and quarantine, a safe and effective vaccine is the only weapon that can potentially offer lasting protection against COVID-19 and stop the current pandemic. According to the WHO, 26 vaccine candidates using various platforms have entered clinical trials as of July 31, 2020 [5]. These candidate vaccines have all been developed in compliance with WHO guidelines that define desired characteristics such as dose regimen, target population, safety, measures of efficacy, and requirements for product stability and storage [6].

Coronaviruses are among the largest known enveloped RNA viruses and cause respiratory illnesses in humans ranging from the common cold to SARS, MERS, as well as the current COVID-19 pandemic [7]. Similar to SARS-CoV, the spike (S) protein of SARS-CoV-2 is the receptor for attachment and cell entry via the cellular receptor hACE2 [1]. Researchers are also adapting antigen design strategies used for SARS-CoV and MERS-CoV spike proteins to develop a candidate vaccine for SARS-CoV-2. A stabilized prefusion form of the MERS spike protein was achieved in 2017 by transferring two proline substitutions between heptad repeat 1 and the central helix analogous to those defined in the HKU1 spike protein. These mutations together with a C-terminus foldon trimerization domain stabilized the spike ectodomain in its prefusion state resulting in a more potent immunogen with dose-sparing properties compared to protein made with the original wild-type sequence [8]. The analogous mutations in the SARS-CoV-2 spike resulted in a homogeneous population of proteins allowing the atomic-level structure to be solved by cryo-electron microscopy [9].

Subunit vaccines such as the spike protein are often poorly immunogenic by themselves and therefore typically require adjuvants to enhance their ability to produce an immune response [10]. Adjuvants can be classified based on their properties into several categories, including aluminum salt-based (aluminum hydroxide and aluminum phosphate), oil emulsion-based such MF59 and AS03, and toll-like receptor (TLR) agonists, including CpG 1018 and monophosphoryl lipid A (AS04) [11]. A previous study employing four different candidate vaccines against SARS-CoV, all of them with and without alum adjuvants successfully elicited neutralizing antibodies and conferred protection against infection; however, tissue damage due to immunopathology was also observed from infiltrating lymphocytes [12]. Historical evidence from animal models suggests, vaccine-primed Th2 and Th17 responses may be associated with immunopathology referred to as vaccine-associated enhanced respiratory disease (VAERD) [13]; therefore, extra caution must be taken in the choice of adjuvants. Synthetic oligodeoxynucleotides with CpG motifs (CpG ODN) are agonists for TLR9 and mimic the activity of naturally occurring CpG motifs found in bacterial DNA. CpG ODN are potent vaccine adjuvants generating Th1-biased responses, thus making them a good choice to mitigate potential Th2 and Th17 induced immunopathology [11]. CpG 1018, a 22-mer CpG ODN containing sequence with a modified phosphorothioate backbone [14], is the adjuvant used in the licensed hepatitis B vaccine HEPLISAV-B^®^, and is the only TLR9 agonist used in an approved vaccine.

In this study, we present data from preclinical studies aimed at developing a COVID-19 candidate subunit vaccine using CHO cell-expressed SARS-CoV-2 S-2P antigen combined with various adjuvants. We have shown that S-2P, when mixed with CpG 1018 and aluminum hydroxide adjuvants, was most effective in inducing antibodies that neutralized pseudovirus and wild-type live virus while minimizing Th2-biased responses with no vaccine-related adverse effects.

## Materials and Methods

### Production of S-2P protein ectodomains from Expi293 and ExpiCHO-S cells

The plasmid expressing SARS-CoV-2 (strain Wuhan-Hu-1 GenBank: MN908947) S protein ectodomain was obtained from Dr. Barney S. Graham (Vaccine Research Center, National Institute of Allergy and Infectious Diseases, USA) and contains a mammalian-codon-optimized gene encoding SARS-CoV-2 S residues 1–1208 with a C-terminal T4 fibritin trimerization domain, an HRV3C cleavage site, an 8×His-tag and a Twin-Strep-tag [9]. The S-2P form was created by mutation of the S1/S2 furin-recognition site 682-RRAR-685 to GSAS to produce a single-chain S0 protein, and the 986-KV-987 was mutated to PP [9].

Expi293 and ExpiCHO-S cells (ThermoFisher) were transfected with the plasmid expressing S-2P protein ectodomains by ExpiFectamine 293 transfection kit and ExpiFectamine CHO transfection kit (ThermoFisher), respectively. The secreted S-2P protein was purified by affinity chromatography. Purification tags were removed by HRV3C protease digestion and the S-2P protein was further purified. The purified S-2P proteins produced from Expi293 and ExpiCHO-S cells were quantified by BCA assay (ThermoFisher), flash frozen in liquid nitrogen and then stored at −80 °C. The ExpiCHO-expressed S-2P were sent to the Electronic Microscopy Laboratory at the Advanced Technology Research Facility (National Cancer Institute) for cryo-EM confirmation.

### Pseudovirus production and titration

To produce SARS-CoV-2 pseudoviruses, a plasmid expressing full-length wild-type Wuhan-Hu-1 strain SARS-CoV-2 spike protein was cotransfected into HEK293T cells with packaging and reporter plasmids pCMVΔ8.91 and pLAS2w.FLuc.Ppuro (RNAi Core, Academia Sinica), using TransIT-LT1 transfection reagent (Mirus Bio). Site-directed mutagenesis was used to generate the D614G variant by changing nucleotide at position 23403 (Wuhan-Hu-1 reference strain) from A to G. Mock pseudoviruses were produced by omitting the p2019-nCoV spike (WT). Seventy-two hours post-transfection, supernatants were collected, filtered, and frozen at −80 °C.The transduction unit (TU) of SARS-CoV-2 pseudotyped lentivirus was estimated by using cell viability assay in response to the limited dilution of lentivirus. In brief, HEK-293T cells stably expressing human ACE2 gene were plated on 96-well plate one day before lentivirus transduction. For the titering of pseudovirus, different amounts of pseudovirus were added into the culture medium containing polybrene. Spin infection was carried out at 1,100 xg in 96-well plate for 30 minutes at 37°C. After incubating cells at 37°C for 16 hr, the culture medium containing virus and polybrene were removed and replaced with fresh complete DMEM containing 2.5 μg/ml puromycin. After treating with puromycin for 48 hrs, the culture media was removed and cell viability was detected by using 10% AlarmaBlue reagents according to manufacturer’s instruction. The survival rate of uninfected cells (without puromycin treatment) was set as 100%. The virus titer (transduction units) was determined by plotting the survival cells versus diluted viral dose.

### Pseudovirus-based neutralization assay

HEK293-hAce2 cells (2×10^4^ cells/well) were seeded in 96-well white isoplates and incubated for overnight. Sera were heated at 56°C for 30 min to inactivate complement and diluted in MEM supplemented with 2 % FBS at an initial dilution factor of 20, and then 2-fold serial dilutions were carried out (for a total of 8 dilution steps to a final dilution of 1:5120). The diluted sera were mixed with an equal volume of pseudovirus (1,000 TU) and incubated at 37 °C for 1 hr before adding to the plates with cells. After the 1-hr incubation, the culture medium was replaced with 50 μL of fresh medium. On the following day, the culture medium was replaced with 100 μL of fresh medium. Cells were lysed at 72 hours post infections and relative luciferase units (RLU) was measured. The luciferase activity was detected by Tecan i-control (Infinite 500). The 50% and 90% inhibition dilution titers (ID_50_ and ID_90_) were calculated considering uninfected cells as 100% neutralization and cells transduced with only virus as 0% neutralization. Reciprocal ID_50_ and ID_90_ geometric mean titers (GMT) were both determined as ID_90_ titers are useful when ID_50_ titer levels are consistently saturating at the upper limit of detection.

### Wild-type SARS-CoV-2 neutralization assay

The neutralization assay with SARS-CoV-2 virus was conducted as previously reported [15]. Vero E6 cells (2.5×10^4^ cells/well) were seeded in 96-well plates and incubated overnight. Sera were heated at 56°C for 30 min to inactivate complement and diluted in serum-free MEM at an initial dilution factor of 20, and then further 2-fold serial dilutions were performed for a total 11 dilution steps to a final dilution of 1:40960. The diluted sera were mixed with an equal volume of SARS-CoV-2 virus at 100 TCID_50_/50 μL (hCoV-19/Taiwan/CGMH-CGU-01/2020, GenBank accession MT192759) and incubated at 37 °C for 2 hr. The sera-virus mixture was then added to 96-well plate with Vero E6 cells and incubated in MEM with 2% FBS at 37 °C for 5 days. After incubation, cells were fixed by adding 4% formalin to each of the wells for 10 min and stained with 0.1% crystal violet for visualization. Results were calculated with the Reed-Muench method for log 50% end point for ID_50_ and log 90% end point for ID_90_ titers.

### Animals

BALB/cJ mice were obtained from the National Laboratory Animal Center, Academia Sinica, Taiwan and BioLASCO Taiwan Co. Ltd. Crl:CD^®^ Sprague Dawley (SD) rats were obtained from BioLASCO Taiwan Co. Ltd. Animal studies were conducted in the Testing Facility for Biological Safety, TFBS Bioscience Inc., Taiwan. All animal work was reviewed and approved by the Institutional Animal Care and Use Committee (IACUC). The Testing Facility’s IACUC animal study protocol approval numbers are TFBS2020-006 and TFBS2020-010.

### Immunization of mice

For antigen formulation, 0.5 mL of SARS-CoV-2 S-2P protein was mixed with either an equal volume of Sigma Adjuvant System S6322 (Sigma), 600 μg/mL Adju-Phos aluminum phosphate (Brenntag), CpG 1018 (200 μg/mL), aluminum hydroxide (1 mg/mL), PBS, or 0.25 mL CpG 1018 (400 μg/mL) plus 0.25 mL aluminum hydroxide (2 mg/mL). Female BALB/cJ mice aged 6–9 weeks were immunized twice at 3 weeks apart as previously described [8]. Total serum anti-S IgG levels were detected with direct ELISA using custom 96-well plates coated with S-2P antigen.

### Cytokine assays

Two weeks after the second injection, mice were euthanized and splenocytes were isolated and stimulated with S-2P protein (2 μg/well) as previously described [16]. For detection of IFN-γ, IL-2, IL-4, and IL-5, the culture supernatant from the 96-well microplates was harvested to analyze the levels of cytokines by ELISA using Mouse IFN-γ Quantikine ELISA Kit, Mouse IL-2 Quantikine ELISA Kit, Mouse IL-4 Quantikine ELISA Kit, and Mouse IL-5 Quantikine ELISA Kit (R&D System). The OD450 values were read by Multiskan GO (ThermoFisher).

### Dose range finding study for single and repeat-dose intramuscular injection (IM) in Sprague Dawley (SD) Rats

To investigate the safety of SARS-CoV-2 S-2P protein adjuvanted with CpG 1018 alone or combined with aluminum hydroxide, pilot toxicity studies were conducted for dose range finding. SD rats aged 6-8 weeks were immunized with 5 μg, 25 μg or 50 μg of S-2P adjuvanted with either CpG 1018 alone or CpG 1018 combined with aluminum hydroxide. The test article or vehicle control was administered intramuscularly to each rat on Day 1 (for single-dose study) and Day 15 (for repeat-dose study). The observation period was 14 days (for single-dose study) and 28 days (for repeat-dose study). Parameters evaluated included clinical signs, local irritation examination, moribundity/mortality, body temperature, body weights, and food consumption during the in-life period. Blood samples were taken for hematology, including coagulation tests and serum chemistry. All animals were euthanized and necropsied for gross lesion examination, organ weights, and histopathology evaluation on injection sites and lungs.

### Statistical analysis

For neutralization assays, geometric mean titers are represented by the heights of bars with 95% confidence intervals represented by the error bars. For cytokine and rat data, heights of bars or symbols represent means with SD represented by error bars.

Dotted lines represent lower and upper limits of detection. Mann-Whitney U-test included in the analysis package in Prism 6.01 (GraphPad) was used to compare between two experimental groups. * = p < 0.05, ** = p < 0.01, *** = p < 0.001

## Results

### Adjuvanted SARS-CoV-2 S-2P induced robust neutralizing antibodies

Sera of BALB/cJ mice vaccinated with HEK293-expressed S-2P with or without adjuvants were assessed using pseudovirus neutralization assays for immunogenicity elicited by S-2P antigen. Reciprocal ID_50_ and ID_90_ geometric mean titers (GMT) were determined as ID_90_ titer is useful when ID_50_ titer levels are consistently saturating at the upper limit of detection. At 1 μg of S-2P, the ID_50_ titers of S-2P alone, with aluminum phosphate, and with Sigma Adjuvant were 259, 2,124, and 5,099, respectively; whereas the ID_90_ titers of the above were 41, 282, and 2007, respectively (Figure S1).

Additional immunogenicity studies were conducted to test the adjuvanted vaccine in different antigen dosages. Similar results were obtained as in the previous experiment (Figure S2). Likewise, both ID_50_ and ID_90_ titers induced by higher antigen dose was stronger than that induced by lower antigen dose. Taken together, these data again confirmed that SARS-CoV-2 S-2P combined with adjuvants induced effective neutralizing antibody, thus indicating early potential and preliminary evidence to pursue development of this candidate COVID-19 vaccine.

### Induction of potent neutralizing antibodies by CpG 1018 and aluminum hydroxide-adjuvanted S-2P

Having established the ability of Expi293-expressed S-2P to induce neutralizing antibodies, we then applied ExpiCHO as the expression system of S-2P antigen for clinical studies and stable clones for commercial production. The S-2P proteins produced in CHO cells and their structure displayed typical spike trimers under cryo-EM (Figure S3), resembling that of Expi293-expressed SARS-CoV-2 S protein, suggesting that CHO cells are feasible in production of S-2P. The above immunogenicity studies showed that the oil in water adjuvant (Sigma adjuvant) vaccinated S-2P could induce effective neutralizing antibody in mice. However, since Sigma adjuvant is not permitted for human therapeutic use and the potential of alum salts in producing Th2-mediated immunopathology, we next examined the potential of Th1-biasing CpG 1018 for clinical use. Aluminum hydroxide was also explored in the following experiment instead of aluminum phosphate as it has been characterized to enhance the potency of CpG adjuvant when used in combination while also retaining the property of inducing Th1 responses [17]. The pseudovirus neutralization assay was performed with sera drawn two weeks after the second injection. At 1 μg of S-2P, the reciprocal ID_50_ GMT of S-2P adjuvanted with CpG 1018, aluminum hydroxide, and with both CpG 1018 and aluminum hydroxide were 245, 3,109, and 5,120, respectively (Figure 1). Similar values were obtained at 5 μg of S-2P (Figure 1). Weaker neutralization titers were observed at 3 weeks after the first injection (Figure S4). Sera from these mice were then examined for the amount of anti-S IgG. CpG 1018 in combination with aluminum hydroxide produced significantly higher titers of anti-S IgG compared to CpG 1018 or aluminum hydroxide alone (Figure 2). The immune sera were further tested for their neutralization capabilities against wild-type SARS-CoV-2 in a neutralization assay. S-2P was able to inhibit SARS-CoV-2 at a concentration of 1 μg, although at lower potency than that of pseudovirus (Figures 1 and 3). The reciprocal ID_50_ GMT of S-2P in the presence of CpG 1018, aluminum hydroxide, and with both CpG 1018 and aluminum hydroxide were approximately 60, 250, and 1,500, respectively (Figure 3). Pseudovirus carrying the current dominant D614G variant spike was also generated and neutralizing antibodies from mice immunized with S-2P with CpG 1018 and aluminum hydroxide were effective against both pseudoviruses carrying the wild-type D614 and mutant D614G versions of spike proteins (Figure 4). Neutralization titers of wild-type virus and pseudovirus and total anti-S IgG titers were all found to be highly correlated with Spearman’s rank correlation coefficients greater than 0.8 (Figure 5).

**Figure 1.**
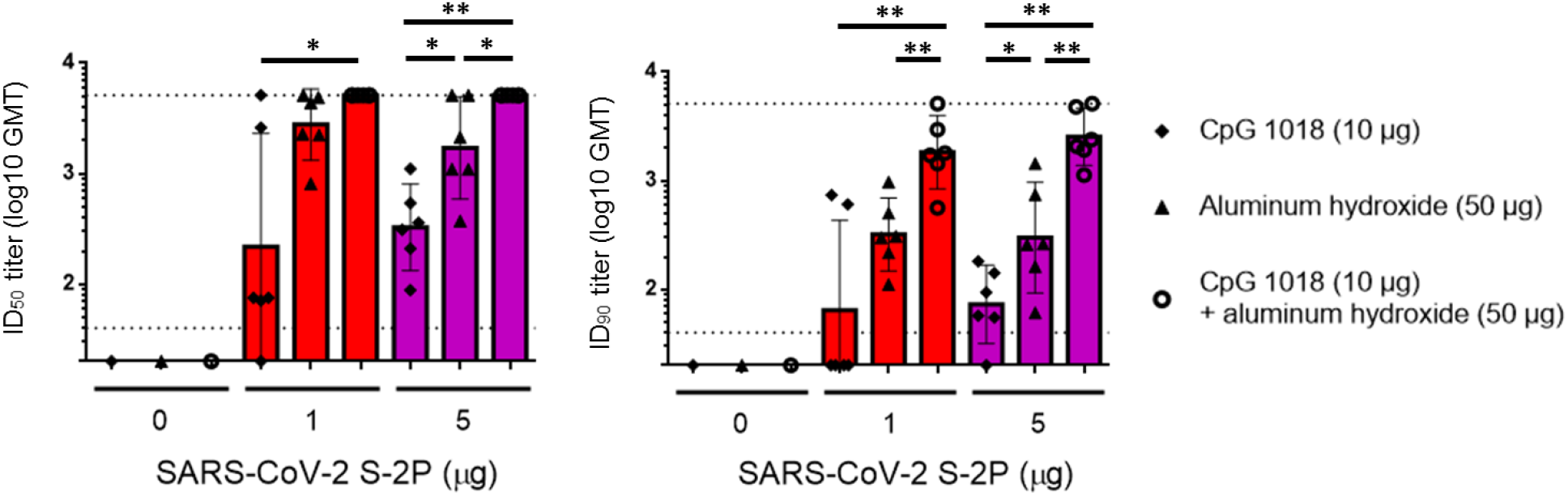
Induction of neutralizing antibodies by CpG 1018 and aluminum hydroxide-adjuvanted SARS-CoV-2 S-2P 2 weeks post-second injection. BALB/c mice were immunized with 2 dose levels of CHO cell-expressed SARS-CoV-2 S-2P adjuvanted with CpG 1018, aluminum hydroxide or combination of both 3 weeks apart and the antisera were harvested at 2 weeks after the second injection. The antisera were subjected to neutralization assay with pseudovirus expressing SARS-CoV-2 spike protein to determine the ID_50_ (left) and ID_90_ (right) titers of neutralization antibodies.

**Figure 2.**
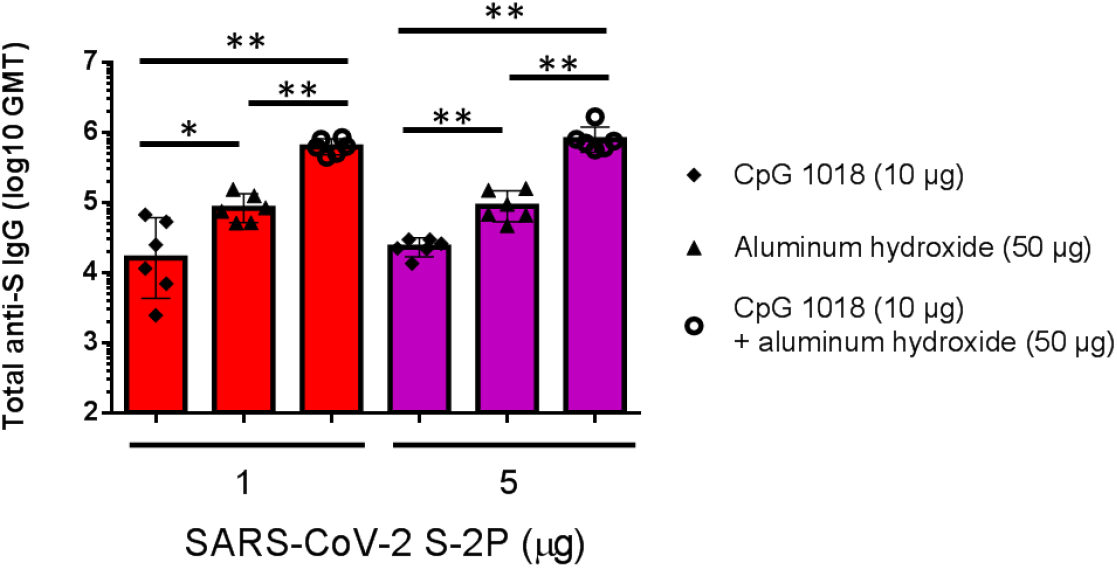
Total anti-S IgG titers in mice immunized with S-2P with adjuvants. Sera from BALB/c mice in Figure 1 immunized with 1 or 5 μg of S-2P with CpG 1018, aluminum hydroxide or combination of both were quantified for the total amount of anti-S IgG with ELISA.

**Figure 3.**
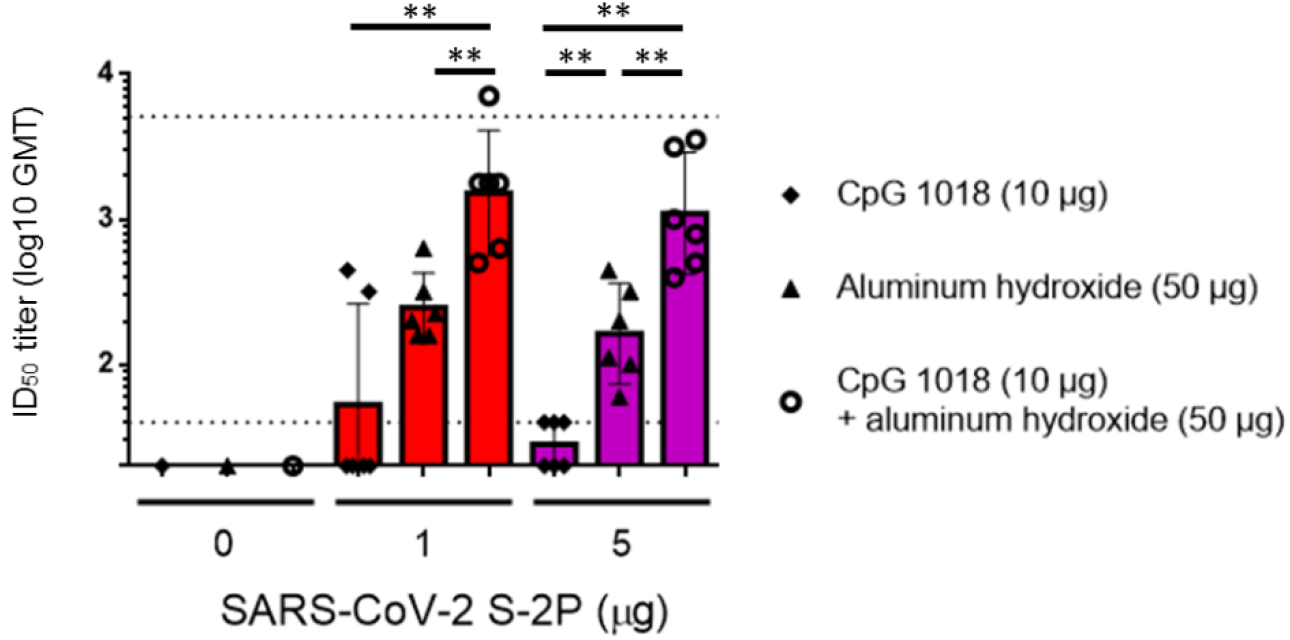
Neutralization of wild-type SARS-CoV-2 virus by antibodies induced by SARS-CoV-2 S-2P adjuvanted with CpG 1018 and aluminum hydroxide. The antisera were collected as described in Figure 2 and subjected to a neutralization assay with wild-type SARS-CoV-2 to determine neutralization antibody titers.

**Figure 4.**
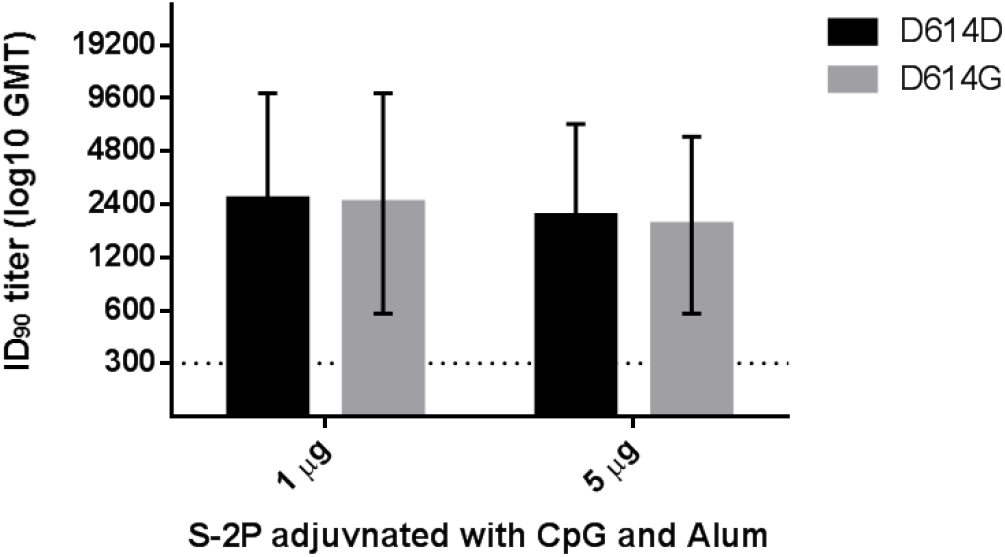
Inhibition of pseudoviruses carrying D614D (wild-type) or D614G (variant) versions of the spike protein by mice immunized with S-2P with CpG 1018 and aluminum hydroxide. The antisera of BALB/c mice immunized with 1 or 5 μg of S-2P with 10 μg CpG 1018 and 50 μg aluminum hydroxide as in Figure 1 were collected. Neutralization assays were performed with pseudoviruses with either D616D or D614G spike proteins.

**Figure 5.**
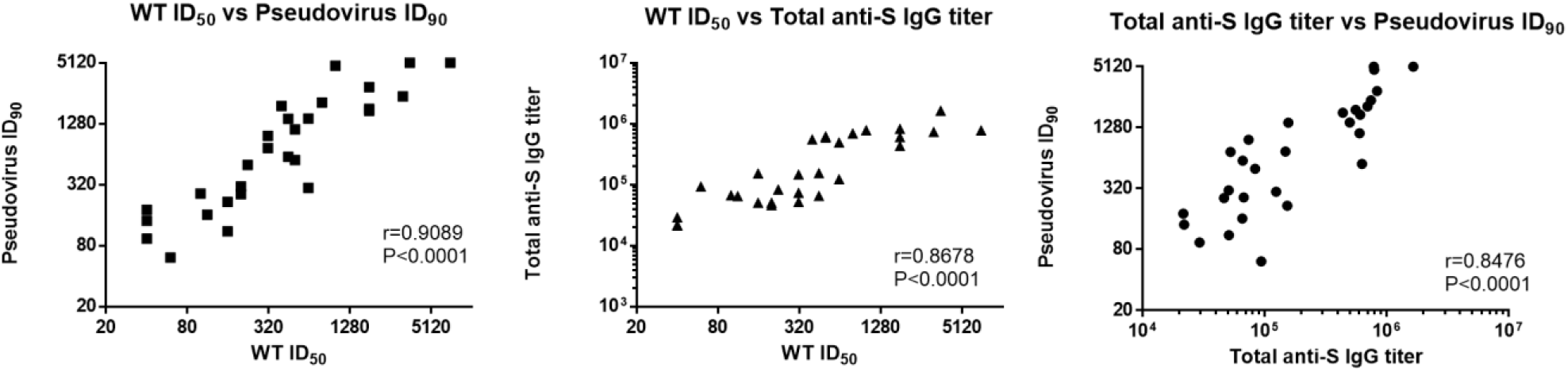
Correlations between SARS-CoV-2 pseudovirus ID_90_, wild-type SARS-CoV-2 ID_50_, and total anti-S IgG titers in mice. Values were tabulated and correlations were calculated with Spearman’s rank correlation coefficient for wild-type SARS-CoV-2 ID_50_ vs pseudovirus ID_90_ (left), wild-type SARS-CoV-2 ID_50_ vs total anti-S IgG titer (middle), and pseudovirus ID_90_ vs total anti-S IgG titer (right).

### CpG 1018 induced Th1 immunity

To identify whether CpG 1018 could induce Th1 responses in our vaccine-adjuvant system, cytokines involved in Th1 and Th2 responses were measured in splenocytes from mice immunized with S-2P with aluminum hydroxide, CpG 1018, or combination of the two. As expected, S-2P adjuvanted with aluminum hydroxide induced limited amounts of IFN-γ and IL-2, the representative cytokines of Th1 response. In contrast, significant increases in IFN-γ and IL-2 were detected most strongly in high antigen dose plus CpG 1018 and aluminum hydroxide (Figure 6). For Th2 response, while the levels of IL-4, IL-5 and IL-6 increased in the presence of aluminum hydroxide and S-2P, addition of CpG 1018 to aluminum hydroxide suppressed the levels of these cytokines (Figure 7). IFN-γ/IL-4, IFN-γ/IL-5, and IFN-γ/IL-6 ratios are strongly indicative of a Th1-biased response and were increased by approximately 36-, 130-, and 2-fold, respectively, in the presence of S-2P combined with CpG 1018 and aluminum hydroxide (Figure 8). These results suggested that the effect of CpG 1018 is dominant over aluminum hydroxide in directing the cell-mediated response towards Th1 response, while retaining high antibody levels.

**Figure 6.**
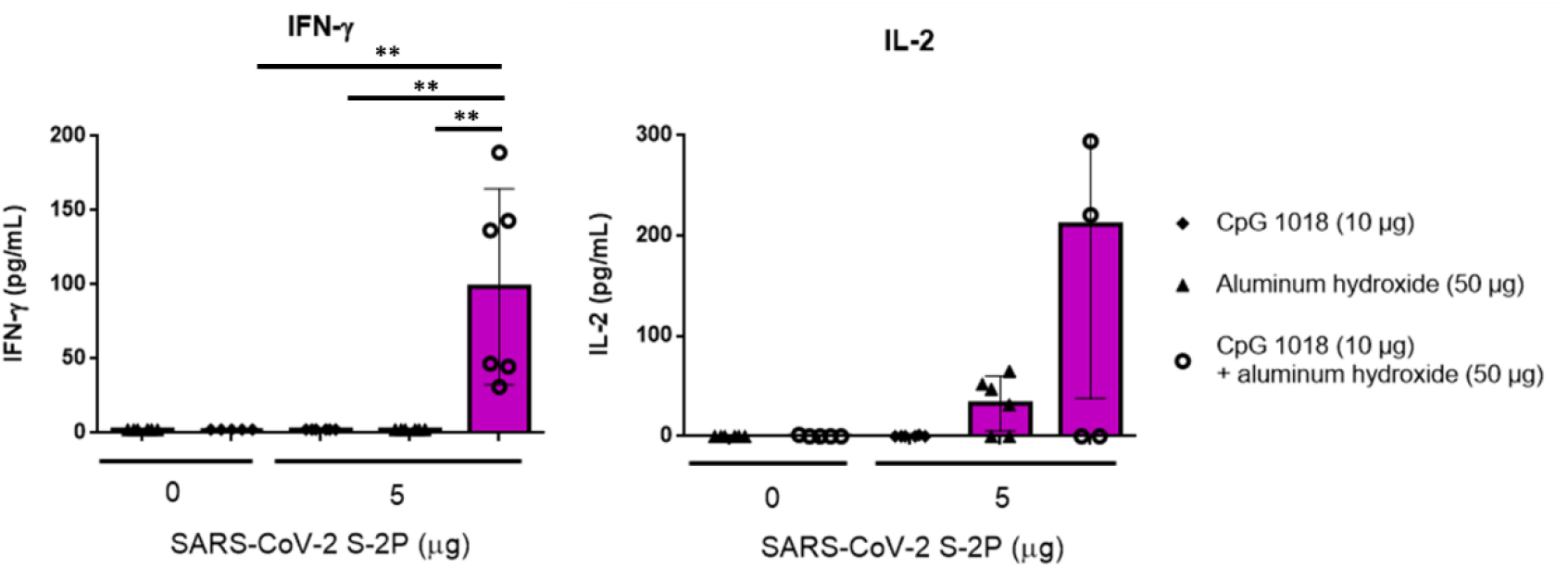
Th1-dependent cytokine production induced by SARS-CoV-2 S-2P adjuvanted with CpG 1018, Aluminum hydroxide, or CpG 1018/Aluminum hydroxide in mice. Two weeks after the second injection, the splenocytes were harvested and incubated with S-2P protein (5 μg), Concanavalin A (0.1 μg; data not shown) for positive control, or complete RPMI 1640 medium only for negative control. After 20 hours incubation, the levels of IFN-γ (left) and IL-2 (right) were analyzed by ELISA.

**Figure 7.**
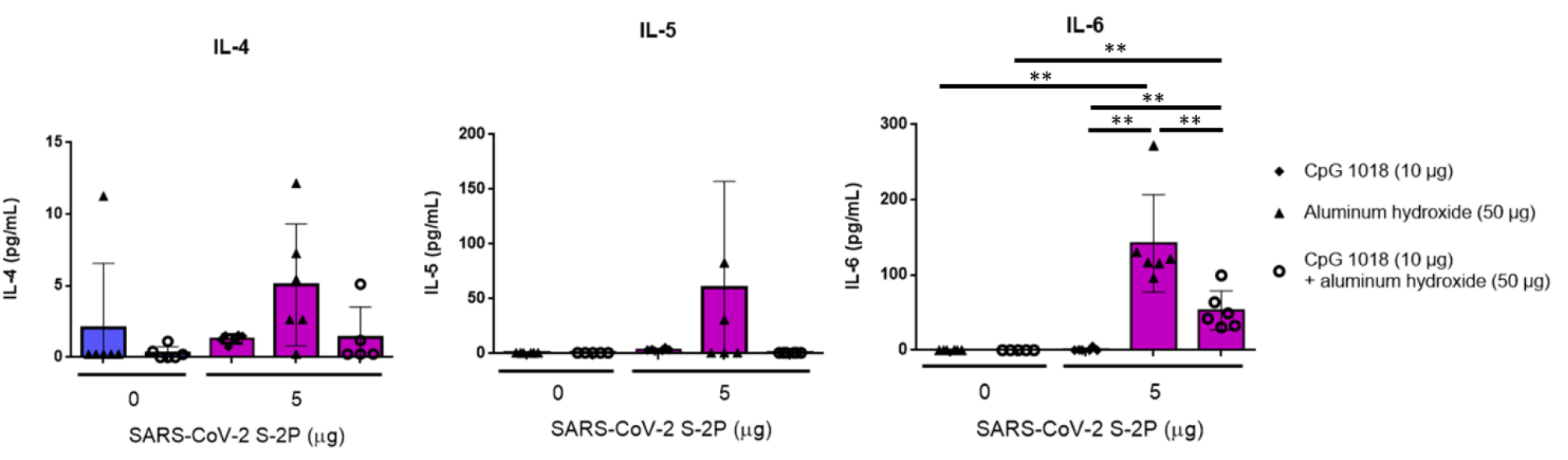
Th2-dependent cytokine production induced by SARS-CoV-2 S-2P adjuvanted with CpG 1018, Aluminum hydroxide, or CpG 1018/Aluminum hydroxide in mice. Two weeks after the second injection, the splenocytes were harvested and stimulated as in Fig. 6. After 20 hours incubation, the levels of IL-4 (left), IL-5 (middle), and IL-6 (right) released from the splenocytes were analyzed. For detection of cytokines, the culture supernatant was harvested to analyze the levels of cytokines by ELISA.

**Figure 8.**
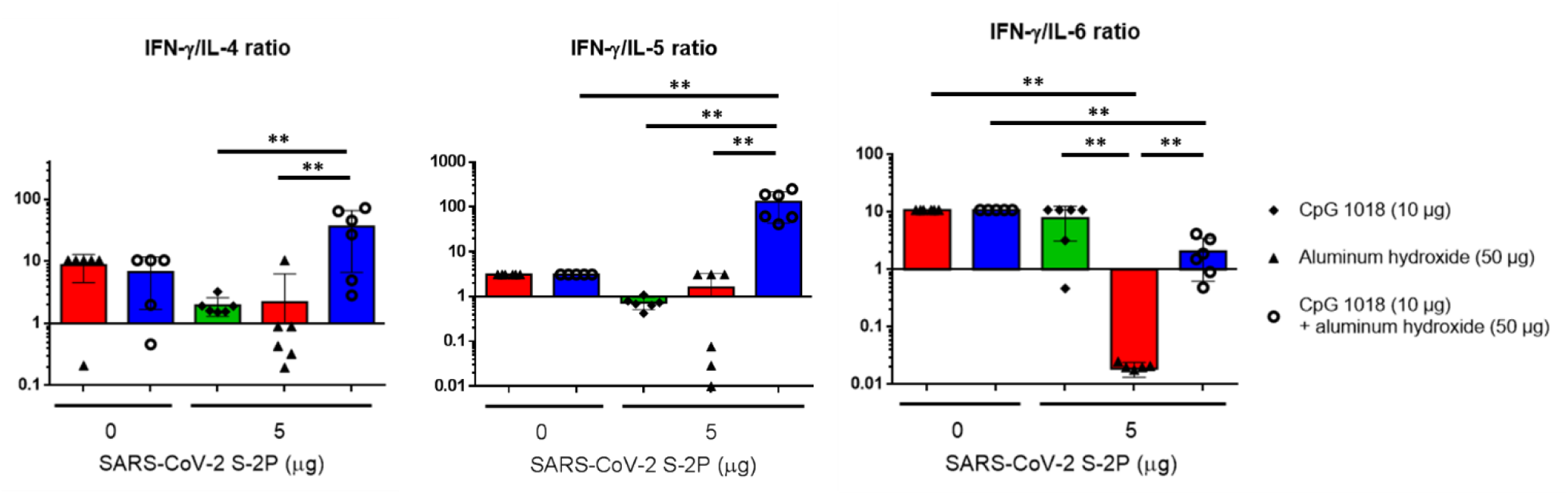
IFN-γ/IL-4, IFN-γ/IL-5, and IFN-γ/IL-6 ratios. IFN-γ, IL-4, IL-5, and IL-6 values from the cytokine assays were used to calculate ratios. Ratio values greater than 1 indicate Th-1 bias whereas ratio less than 1 indicate Th-2 bias responses.

### S-2P did not result in systemic adverse effects in rats

To elucidate the safety and potential toxicity of the vaccine candidate, 5 μg, 25 μg or 50 μg of S-2P adjuvanted with CpG 1018 or CpG 1018 combined aluminum hydroxide were administered to SD rats for single-dose and repeat-dose studies. No mortality, abnormality of clinical signs, differences in body weight changes, body temperature, nor food consumption were observed in either gender that could be attributed to S-2P (with or without adjuvant) with single dose administration (Figures 9 and 10). Increased body temperature at 4-hr or 24-hr after dosing was found in both genders of single-dose study and repeat-dose study; however, these temperature changes were moderate and were recovered after 48-hr in both genders of all treated groups including controls (PBS) (Figure 9). No gross lesions were observed in organs of most of the male and female rats with single-dose and two-dose administration, except for one male rat which was deemed to be non-vaccine-related. In conclusion, S-2P protein, with CpG 1018 or CpG 1018 + aluminum hydroxide as adjuvants administrated intramuscularly once or twice to SD rats did not induce any systemic adverse effect.

**Figure 9.**
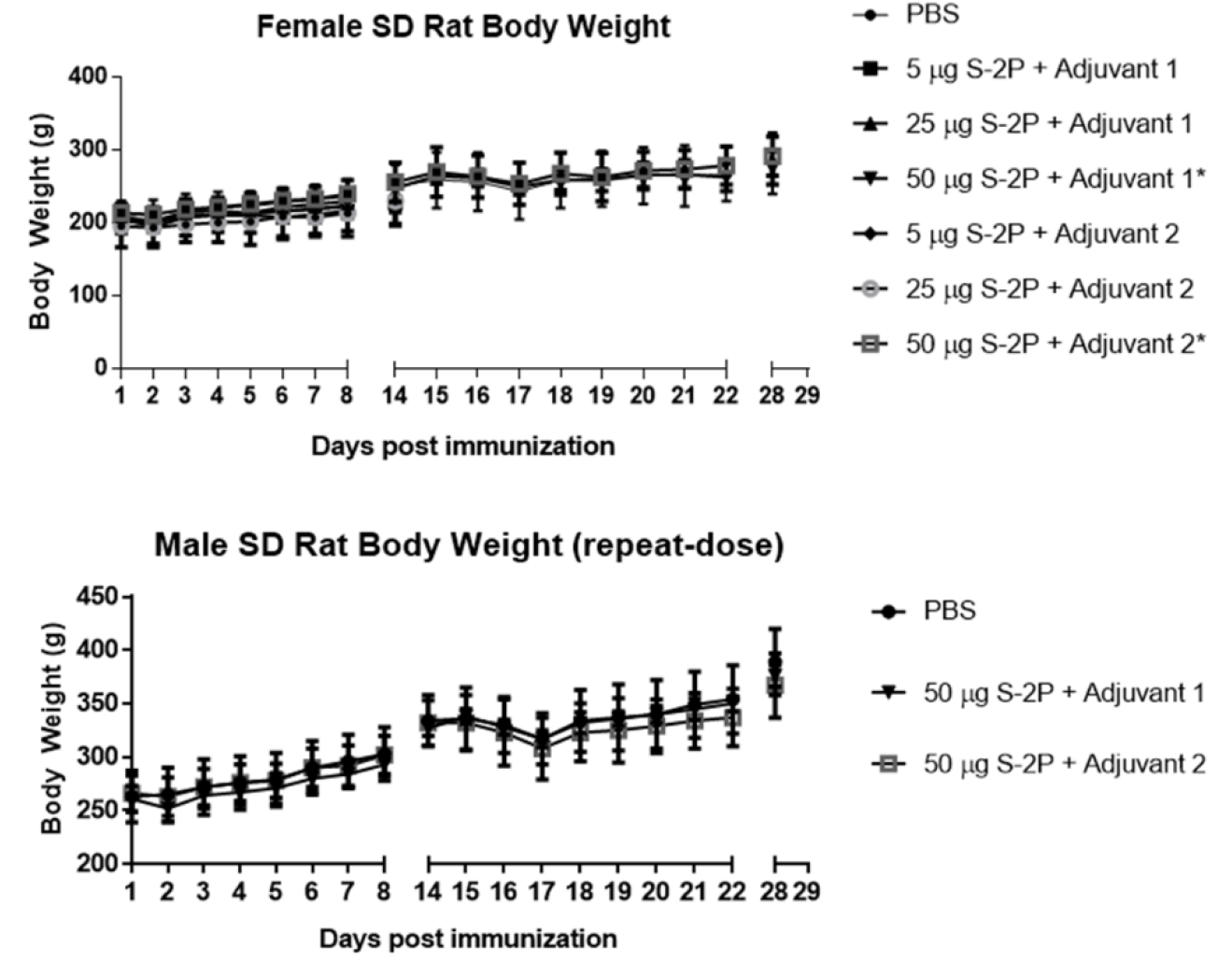
Female (top) and male (bottom) body weight of SD rats immunized with indicated amount of S-2P with adjuvants. Adjuvant 1 = CpG 1018, Adjuvant 2 = CpG 1018 + aluminum hydroxide.

**Figure 10.**
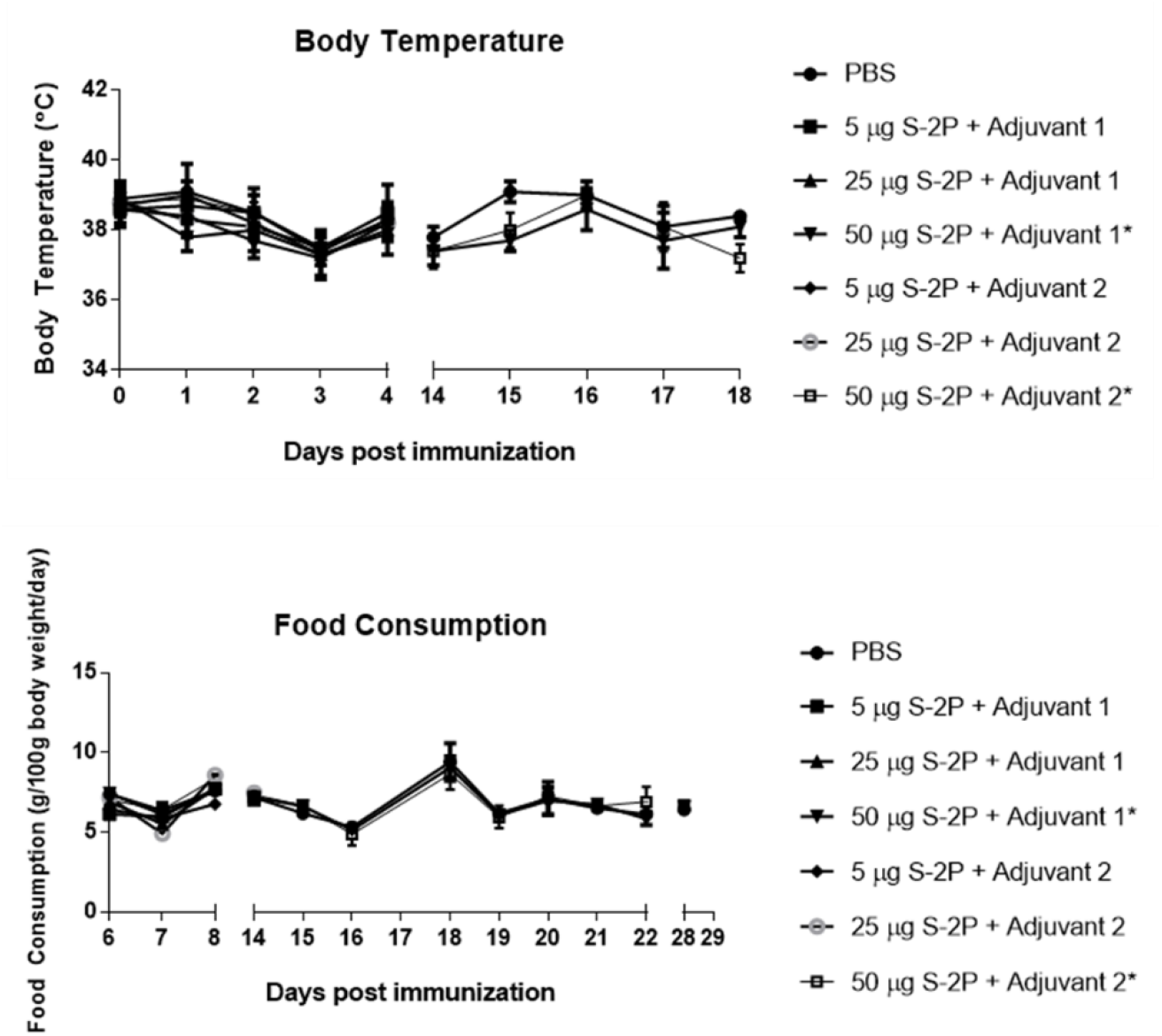
Body temperature (top) and food consumption (bottom) of female and male SD rats immunized with indicated amount of S-2P with adjuvants. Adjuvant 1 = CpG 1018, Adjuvant 2 = CpG 1018 + aluminum hydroxide.

## Discussion

In this study, we showed that in mice, two injections of a subunit vaccine consisting of the prefusion spike protein (S-2P) adjuvanted with CpG 1018 and aluminum hydroxide was effective in inducing potent neutralization activity against both pseudovirus expressing wild-type and D614G variant spike proteins, and wild-type SARS-CoV-2. The combination of S-2P with CpG 1018 and aluminum hydroxide elicited Th1-dominant immune responses with high neutralizing antibody levels in mice and showed no major adverse effects in rats. We also successfully scaled-up yield of S-2P by establishing stable CHO cell clones expressing S-2P protein and improved the purification process at a sufficient quantity of antigen for the production of a commercial vaccine. Spike is a highly glycosylated protein and we chose CHO-cell production to achieve mammalian glycosylation patterns that will include complex glycans and may be important for immunogenicity. Although the leading subunit protein COVID-19 vaccines by developers such as Sanofi Pasteur and Novavax are made in baculovirus, the insect cell produce protein with man-9 glycosylation that may be sufficient for immune response induction, but may not recapitulate the antigenicity of virus grown in mammalian cells [5, 18–19]. Animal challenge studies will be conducted at a future date to examine the safety and efficacy of our candidate vaccine. Based on our results and in accordance with the International Coalition of Medicines Regulatory Authorities (ICMRA), we plan to move forward with first-in-human clinical trials and conduct preclinical studies in parallel to expedite vaccine development in the current COVID-19 pandemic.

We have successfully shown robust immunogenicity elicited by adjuvanted SARS-CoV-2 S-2 (Figures 1, 2, S1, and S2). Much stronger neutralizing antibody responses were detected in mice when 1 μg or 5 μg of S-2P protein was adjuvanted with 10 μg of CpG 1018 and 50 μg of aluminum hydroxide than with either adjuvant alone (Figure 1). S-2P in conjunction with CpG 1018 and aluminum hydroxide induced potent anti-S antibodies that were effective against wild-type virus (Figures 2 and 3). We have shown that high degrees of correlation between neutralization titers of pseudovirus, wild-type virus, and anti-S IgG titers (Figure 5), raising the potential that anti-S IgG titer could be used as surrogate for vaccine potency instead of performing pseudovirus or wild-type virus assays. During the course of vaccine development against fast-evolving RNA viruses such as SARS-CoV-2, it is important that the protection offered by the vaccine could extend to variants that could otherwise drastically reduce the effectiveness of neutralizing antibody. Strains harboring the D614G mutation in the spike protein were first observed in Europe in February 2020 and overtime has become the global dominant variant [20]. Our results of the pseudovirus neutralization assay showed cross-reaction of these antibodies with the dominant circulating stain D614G with similar titer levels (Figure 4). Therefore, we confirmed that S-2P was able to generate antibodies effective against both the original wild-type strain and its variant. Neutralization titers of antibodies against different strains of wild-type viruses should be investigated in the future, but our results indicate the potential of this candidate vaccine to provide broad spectrum protection against COVID-19 infection.

Although moderate IL-4 production was detected in mice receiving 5 μg of S-2P combined with CpG 1018 and aluminum hydroxide, the IFN-γ/IL-4 ratio was 16-fold higher than those receiving 5 μg of S-2P adjuvanted with aluminum hydroxide alone. These results suggested that CpG 1018, even in the presence of aluminum hydroxide could steer the immune response away from Th2 to a Th1 response. Moreover, these mice produce a limited amount of IL-5, which is a key mediator in eosinophil activation and major regulator of eosinophil accumulation in tissues [21]. Previous studies showed that the lung-infiltrating eosinophils were a common indication of Th2-biased immune responses seen in animal models testing SARS-CoV vaccine candidates [22]. The finding that IL-5 production was inhibited by the S-2P adjuvanted with CpG 1018 plus aluminum hydroxide suggests that it would be less likely to induce immune responses resulting in eosinophil infiltration in lung. Th1 and Th2-biased responses are determined by factors, including administration routes, antigen and adjuvant characteristics, and cytokines [11]. Our results showed that S-2P per se is unlikely to skew the immune response towards Th1, but in the presence of an adjuvant such as CpG 1018, S-2P can direct the immune response towards Th1. Thus, we have shown that S-2P adjuvanted with CpG 1018 plus aluminum hydroxide is a potential formulation for COVID-19 vaccine development.

Single-dose or repeat-dose administration of S-2P protein adjuvanted with CpG 1018 and aluminum hydroxide was well tolerated in rats in both genders, supporting human clinical trials in young healthy adults. GLP toxicology study of S-2P in combination of higher dose of CpG 1018 and aluminum hydroxide will be conducted to explore safety of the formulated S-2P for dose escalation study in human clinical trials with the elderly and those with chronic health conditions such as diabetes and cardiovascular diseases, who may require a higher adjuvant dose to boost the immune systems. As a two-dose regimen of S-2P formulated with CpG 1018 and aluminum hydroxide induced potent neutralizing activity, our future plans will include testing single-dose regimens.

Our study showed that CHO-derived S-2P proteins elicited robust immune responses in mice, indicating that CHO cell is an appropriate platform for stable S-2P production in vaccine development. Other vaccines using CHO cells to produce antigens include hepatitis B vaccines GenHevacB and Sci-B-Vac [23]. To this date, we have established stable CHO cell clones expressing S-2P and the one with the highest yield will be selected to produce master cell bank for large scale GMP production of commercial vaccine.

The rapid spread of SARS-CoV-2 and urgent need for an effective vaccine call for its development using readily available and proven technologies. The spike protein is the main receptor binding and membrane fusion protein, which serves as the major antigen target for COVID-19 vaccine development. We have demonstrated in this study that the S-2P combined with the advanced adjuvant CpG 1018, the adjuvant contained in the FDA-approved adult hepatitis B vaccine (HEPLISAV-B), in combination with aluminum hydroxide induced potent Th1-biased immune responses to prevent wild-type virus infections while retaining high antibody levels that show cross-neutralization of variant viruses. Therefore, this vaccine formulation serves as an ideal vaccine candidate in alleviating the burden of the global COVID-19 pandemic.

## Acknowledgements

We thank Dr. Barney S. Graham, Dr. Kizzmekia S. Corbett and members of the Graham lab for providing the S-2P plasmid and initial batch of S-2P protein, as well as providing technical guidance and helpful advices. We are grateful for the participation of Dr. Michael D. Malison, Dr. Han van den Bosch, Dr. Ya-Shan Chuang, and Yu-Fan Tu for manuscript preparation and constructive comments. In addition, we would like to acknowledge Dr. Yu-Chi Chou for the pseudovirus neutralization assay at the RNAi Core Facility, Academia Sinica. We also thank team members at TFBS Bioscience Incorporation for animal experiments and cytokine detection.

---

**Figure S1.**
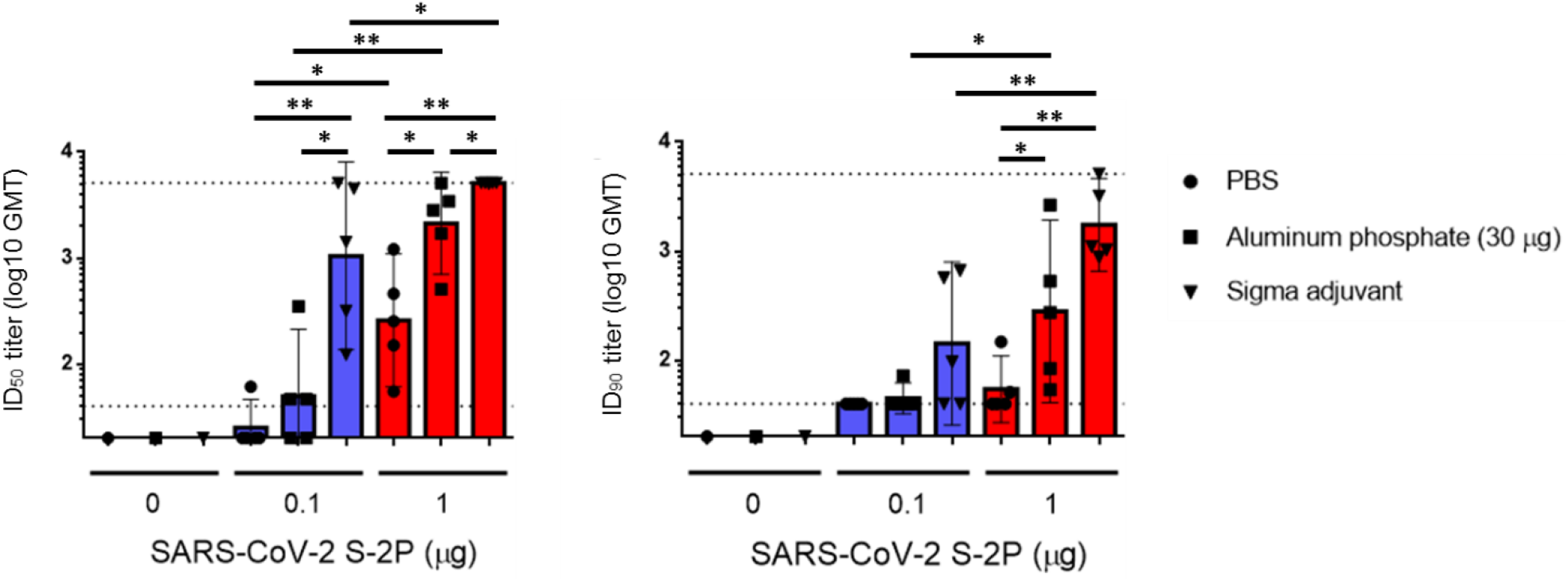
Immunogenicity elicited by adjuvanted SARS-CoV-2 S-2P. BALB/c mice were immunized with 2 injections of HEK 293 cell-expressed SARS-CoV-2 S-2P alone or combined with various adjuvants 3 weeks apart and the antisera were harvested at 2 weeks after the second injection. The antisera were subjected to a neutralization assay with pseudovirus expressing SARS-CoV-2 spike protein to determine the ID_50_ (left) and ID_90_ (right) titers of neutralization antibodies.

**Figure S2.**
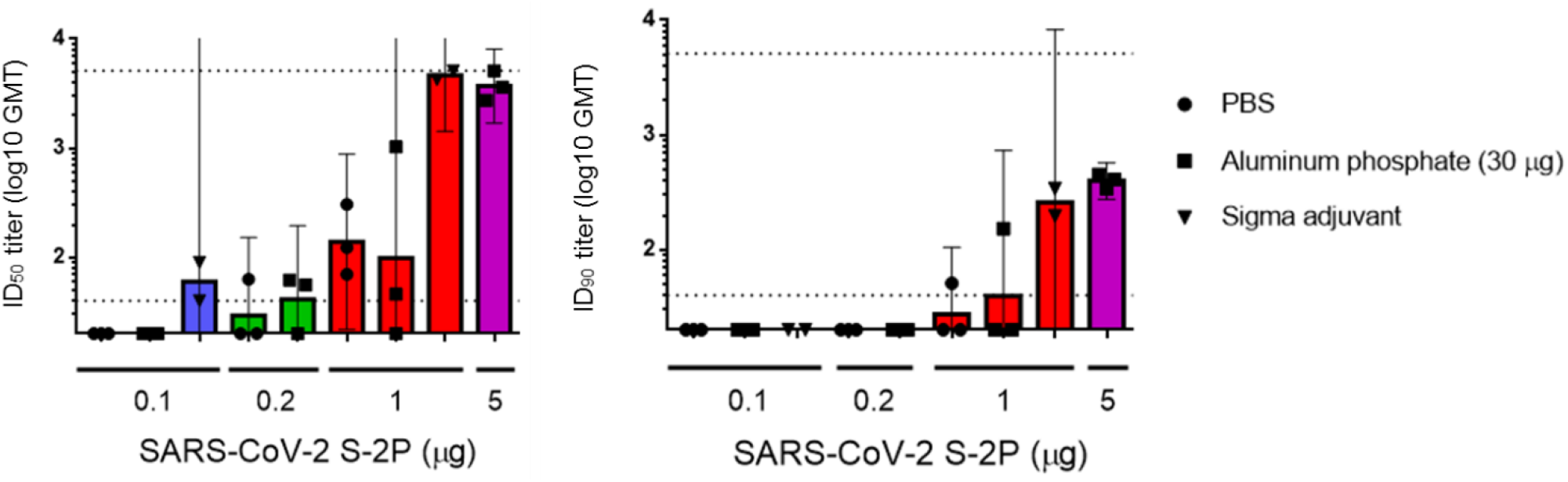
Neutralization antibodies induced by adjuvanted SARS-CoV-2 S-2P. BALB/c mice were immunized with different dose levels of Expi293 cell-expressed SARS-CoV-2 S-2P protein as described in Figure 1. 2 weeks after the second injection, the antisera were subjected to neutralization assay with pseudovirus expressing SARS-CoV-2 spike protein to determine the ID_50_ (left) and ID_90_ (right) titers of neutralization antibodies.

**Figure S3.**
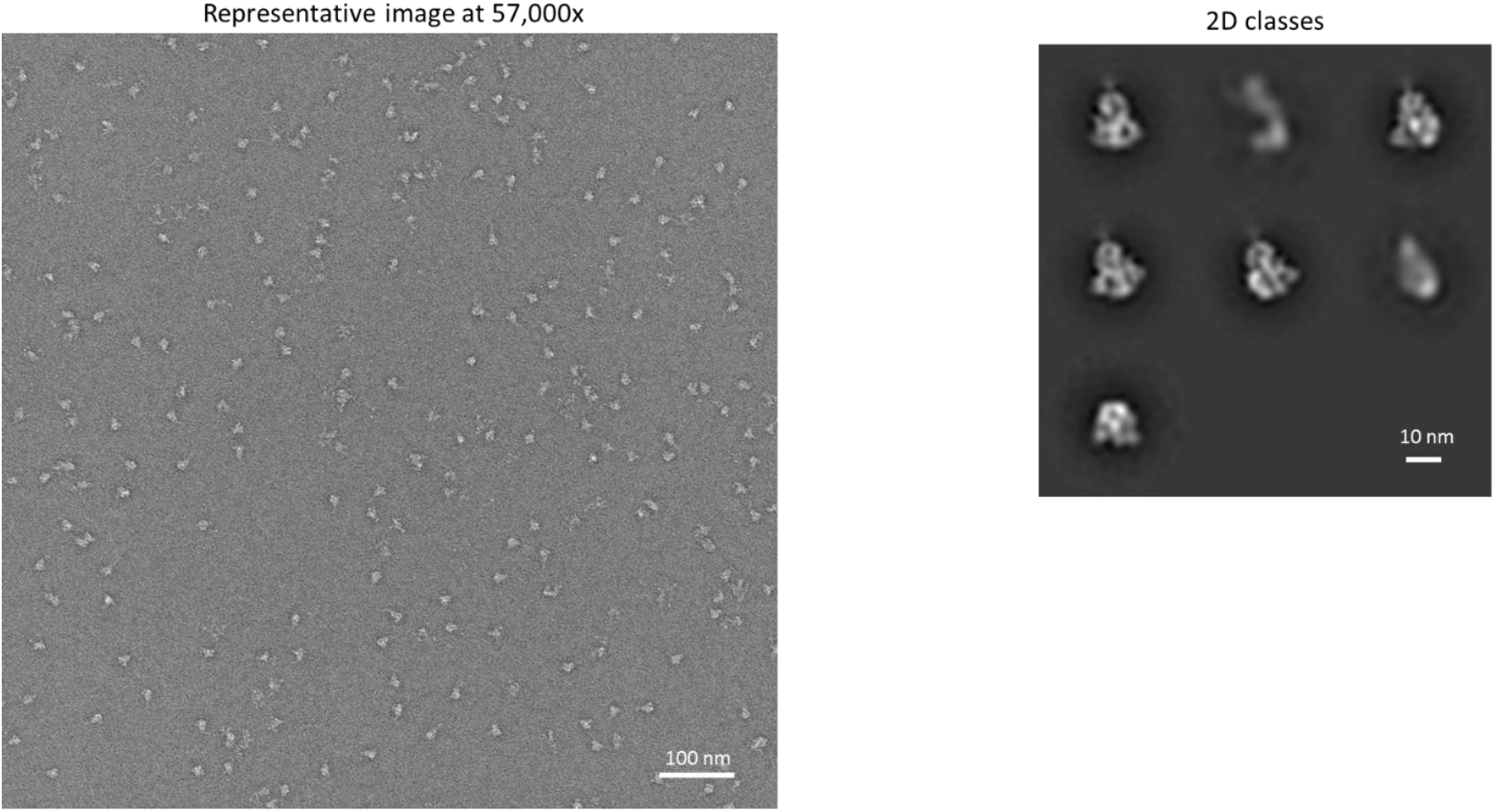
Assembled spike trimers of SARS-CoV2 S-2P under EM (left) and corresponding to ordered spike molecules 2D classes (right). SARS-CoV2 S-2P was transiently expressed by ExpiCHO cells. The sample contains primarily assembled spike trimers. Most particles contributed to 2D classes corresponding to ordered spike molecules.

**Figure S4.**
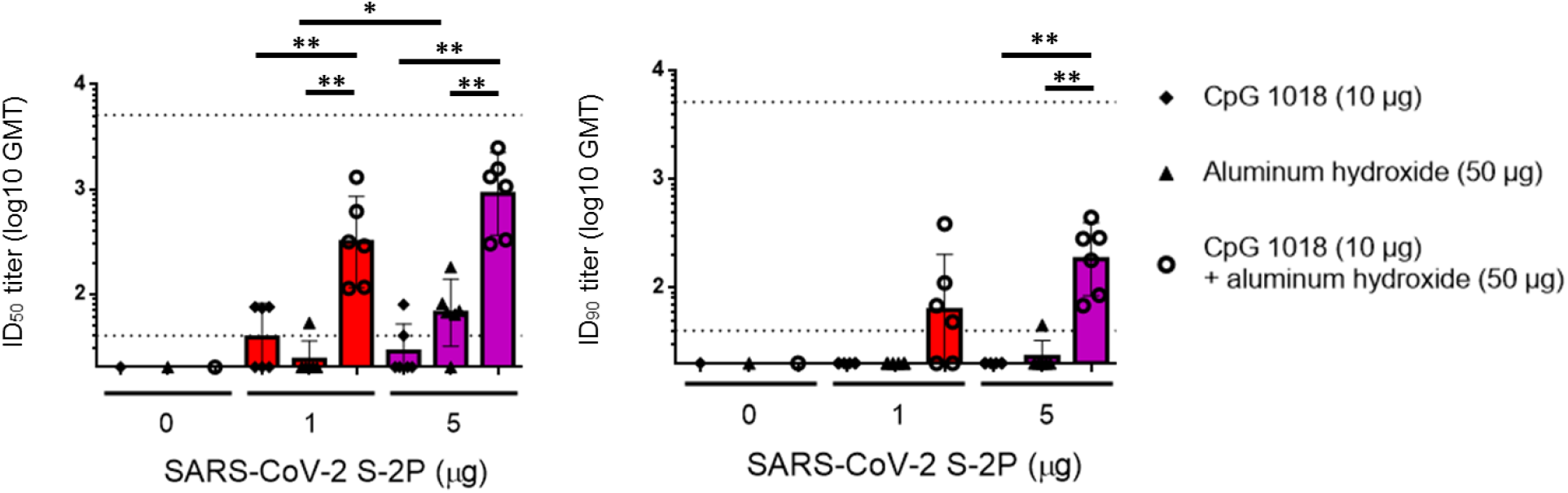
Neutralizing antibody responses in BALB/c mice 3 weeks after first injection of CpG 1018 and aluminum hydroxide-adjuvanted SARS-CoV-2 S-2P. BALB/c mice were immunized with 2 injections of CHO cell-expressed SARS-CoV-2 S-2P adjuvanted with CpG 1018, aluminum hydroxide or combination of both 3 weeks apart and the antisera were harvested at 3 weeks after the first injection. The antisera were subjected to neutralization assay with pseudovirus expressing SARS-CoV-2 spike protein to determine the ID_50_ (left) and ID_90_ (right) titers of neutralization antibodies.

## References

1. Zhou P, Yang XL, Wang XG, Hu B, Zhang L, Zhang W, Si HR, Zhu Y, Li B, Huang CL, Chen HD. Discovery of a novel coronavirus associated with the recent pneumonia outbreak in humans and its potential bat origin. BioRxiv. 2020 Jan 1.

2. ArcGIS Dashboards. Gisanddata.maps.arcgis.com. 2020 [retrieved 05 August 2020]. Available from: https://gisanddata.maps.arcgis.com/apps/opsdashboard/index.html#/bda7594740fd40299423467b48e9ecf6

3. Gates B. Responding to Covid-19—a once-in-a-century pandemic?. New England Journal of Medicine. 2020 Apr 30;382(18):1677-9.

4. Schäferhoff M, Yamey G, McDade K. Funding the development and manufacturing of COVID-19 vaccines: The need for global collective action. Brookings. 2020 [retrieved 1 June 2020]. Available from: https://www.brookings.edu/blog/future-development/2020/04/24/funding-the-development-and-manufacturing-of-covid-19-vaccines-the-need-for-global-collective-action/

5. WHO R&D Blueprint. DRAFT Landscape of COVID-19 Candidate Vaccines – 31 July 2020. Available from: https://www.who.int/who-documents-detail/draft-landscape-of-covid-19-candidate-vaccines

6. WHO R&D Blueprint. Target Product Profiles for COVID-19 Vaccines – 17 April 2020. Available from: https://www.who.int/who-documents-detail/who-working-group-target-product-profiles-for-covid-19-vaccines

7. Gorbalenya AE. Severe acute respiratory syndrome-related coronavirus–The species and its viruses, a statement of the Coronavirus Study Group. BioRxiv. 2020 Jan 1.

8. Pallesen J, Wang N, Corbett KS, Wrapp D, Kirchdoerfer RN, Turner HL, Cottrell CA, Becker MM, Wang L, Shi W, Kong WP. Immunogenicity and structures of a rationally designed prefusion MERS-CoV spike antigen. Proceedings of the National Academy of Sciences. 2017 Aug 29;114(35):E7348–57.

9. Wrapp D, Wang N, Corbett KS, Goldsmith JA, Hsieh CL, Abiona O, Graham BS, McLellan JS. Cryo-EM structure of the 2019-nCoV spike in the prefusion conformation. Science. 2020 Mar 13;367(6483):1260–3.

10. Lee S, Nguyen MT. Recent advances of vaccine adjuvants for infectious diseases. Immune network. 2015 Apr 1;15(2):51–7.

11. Shi S, Zhu H, Xia X, Liang Z, Ma X, Sun B. Vaccine adjuvants: Understanding the structure and mechanism of adjuvanticity. Vaccine. 2019 May 27;37(24):3167–78.

12. Tseng CT, Sbrana E, Iwata-Yoshikawa N, Newman PC, Garron T, Atmar RL, Peters CJ, Couch RB. Immunization with SARS coronavirus vaccines leads to pulmonary immunopathology on challenge with the SARS virus. PloS one. 2012 Apr 20;7(4):e35421.

13. Hotez PJ, Corry DB, Bottazzi ME. COVID-19 vaccine design: the Janus face of immune enhancement. Nature Reviews Immunology. 2020 Jun;20(6):347–8.

14. Campbell JD. Development of the CpG adjuvant 1018: a case study. In Vaccine Adjuvants 2017 (pp. 15–27). Humana Press, New York, NY.

15. Huang CG, Lee KM, Hsiao MJ, Yang SL, Huang PN, Gong YN, Hsieh TH, Huang PW, Lin YJ, Liu YC, Tsao KC. Culture-based virus isolation to evaluate potential infectivity of clinical specimens tested for COVID-19. Journal of Clinical Microbiology. 2020 Jun 9.

16. Lu B, Huang Y, Huang L, Li B, Zheng Z, Chen Z, Chen J, Hu Q, Wang H. Effect of mucosal and systemic immunization with virus-like particles of severe acute respiratory syndrome coronavirus in mice. Immunology. 2010 Jun;130(2):254–61.

17. Thomas LJ, Hammond RA, Forsberg EM, Geoghegan-Barek KM, Karalius BH, Marsh, Jr HC, Rittershaus CW. Co-administration of a CpG adjuvant (VaxImmuneTM, CPG 7909) with CETP vaccines increased immunogenicity in rabbits and mice. Human vaccines. 2009 Feb 1;5(2):79–84.

18. Tian JH, Patel N, Haupt R, Zhou H, Weston S, Hammond H, Lague J, Portnoff AD, Norton J, Guebre-Xabier M, Zhou B. SARS-CoV-2 spike glycoprotein vaccine candidate NVX-CoV2373 elicits immunogenicity in baboons and protection in mice. bioRxiv. 2020 Jan 1.

19. O’Flaherty R, Bergin A, Flampouri E, Mota LM, Obaidi I, Quigley A, Xie Y, Butler M. Mammalian cell culture for production of recombinant proteins: A review of the critical steps in their biomanufacturing. Biotechnology Advances. 2020 May 13:107552.

20. Korber B, Fischer WM, Gnanakaran S, Yoon H, Theiler J, Abfalterer W, Hengartner N, Giorgi EE, Bhattacharya T, Foley B, Hastie KM. Tracking changes in SARS-CoV-2 Spike: evidence that D614G increases infectivity of the COVID-19 virus. Cell. 2020 Jul 3.

21. Lopez AF, Begley CG, Williamson DJ, Warren DJ, Vadas MA, Sanderson CJ (May 1986). “Murine eosinophil differentiation factor. An eosinophil-specific colony-stimulating factor with activity for human cells”. J. Exp. Med. 163 (5): 1085–1099.

22. Tseng CT, Sbrana E, Iwata-Yoshikawa N, et al. Immunization with SARS coronavirus vaccines leads to pulmonary immunopathology on challenge with the SARS virus [published correction appears in PLoS One. 2012;7(8). doi:10.1371/annotation/2965cfae-b77d-4014-8b7b-236e01a35492]. PLoS One. 2012;7(4):e35421. doi:10.1371/journal.pone.0035421

23. Perazzo P, Valle NRD, Sordelli A, Gonzalez RH, Cuestas ML, et al. (2015) Nanotechnology, Drug Delivery Systems and their Potential Applications in Hepatitis B Vaccines. Int J Vaccines Vaccin 1(2): 00007.

